# Three-dimensional Isotropic Imaging of Live Suspension Cells Enabled by Droplet Microvortices

**DOI:** 10.1101/2023.12.01.569311

**Authors:** Braulio Cardenas-Benitez, Richard Hurtado, Xuhao Luo, Abraham P. Lee

## Abstract

Three-dimensional (3D) imaging of non-adherent cells in suspension media is challenging due to their propensity to drift when not fixed to a substrate, as required by optical sectioning technologies. Resolution differences in the lateral versus depth directions typically present in those systems further complicates single-cell morphometry of cellular features indicative of effector functions, such as cytosol and organelle volumetric distribution, and cell membrane topography. Here, we present a method for 3D fluorescent isotropic imaging of live, non-adherent single cells encapsulated in picoliter droplets using Optical Projection Tomography (OPT) enabled by droplet microvortices. Our microfluidic platform features a droplet trap array that leverages flow-induced droplet interfacial shear to generate intra-droplet microvortices, which in turn are modulated to rotate single-cells on their axis to enable OPT-based imaging. This strategy allows observation of cells encapsulated inside non-toxic isotonic buffer droplets and facilitates scalable OPT acquisition by the simultaneous spinning of hundreds of cells. Specifically, we demonstrate 3D imaging of live myeloid and lymphoid cells in suspension, including K562 cells, as well as naïve and activated T cells—small cells prone to movement in their suspended phenotype. In addition, morphometry of primary T cells under different immunological activation states allowed us to identify six distinct nuclear content distributions, which differ from the conventional 2D images depicting spheroid and bean-like nuclear shapes commonly associated with lymphocytes. This Arrayed-Droplet Optical Projection Tomography (ADOPT) technology is capable of isotropic, single live-cell 3D imaging and has the potential to perform large-scale morphometry of immune cell effector function states, while providing compatibility with microfluidic droplet operations.

## Introduction

Understanding the three-dimensional (3D) topography of cells and their intracellular structures plays a central role in biology and biomedical research. Many pivotal discoveries in immune cell biology, such as elucidating the structural organization of the immunological synapse in T cells^1^ or the spatial visualization of intracellular components like the 3D nuclear distribution of cells,^2^ have resulted from advances in optical sectioning technologies.^3^ Despite these advances, there exist key technical challenges inherent to live cell 3D optical imaging, including diffraction-limited resolution and point spread function (PSF) anisotropy, depth-dependent signal intensity degradation, photobleaching, phototoxicity, the cost and complexity of optics, and spatiotemporal resolution capable of resolving unfixed cell motion. In particular, imaging non-adherent cells in 3D, like immune system lymphocytes, poses a unique challenge for optical sectioning methods due to their tendency to drift when not bound to a substrate. This critically impacts live cell topographic and intracellular morphometric measurements in their suspended phenotype,^4^ which ideally requires isotropic and depth-independent signal acquisition over the cell volume while minimizing cell motion artifacts.

Confocal laser scanning microscopy (CLSM) continues to be the method of choice among biologists for performing 3D live cell imaging, owing to its optical sectioning capabilities, its versatility for imaging a wide variety of fluorescence dyes and markers, and enhanced signal-to-noise ratio (SNR) via out-of-focus light rejection.^3^ The method, however, suffers from poor temporal resolution, volumetric bleaching, and significant depth-dependent signal degradation even across the volume of a single cell.^5^ Structured illumination microcopy (SIM) and lattice light-sheet microscopy (LLSM) offer significantly higher spatio-temporal resolution, albeit at increased instrumentation complexity and cost, low-throughput for population analysis, and inability to image suspended, non-adherent single-cells without treatment of the surface imaging substrate.^6^ In contrast, 3D imaging flow cytometry (IFC) employing light-sheet illumination,^6^ and tomographic imaging flow cytometry (tIFC)^7^ techniques have addressed throughput limitations, but have in practice demonstrated limited optical resolutions (∼0.5-1 µm, although slower flow rates have been proposed to reach the optical limit^6^). Furthermore, the complex illumination schemes required in those techniques require optical components that can be prohibitively expensive for some research laboratories.

Previous techniques based on Optical Projection Tomography (OPT) have been proposed to image single-cells in their suspended phenotype with spatial isotropic resolution.^8^ OPT is an imaging method that utilizes a collection of 2D pseudo projections of an object taken at varying angular orientations to reconstruct its 3D structure.^4^ The method was originally devised to analyze millimeter-sized specimens,^9^ and has been adapted to study cell topography as well as intracellular phenomena. In epifluorescence Live Cell Computed Tomography (LCCT), 3D reconstructions based on optical projection tomography (OPT) of suspension cell lines, such as leukemic K562 cells, allowed for isotropic quantification of mitochondrial and nuclear dynamics.^4^ Highly inclined and laminated fluorescence computed tomography (HILO-fCT), another variation of LCCT, offers reduced photobleaching and improved image contrast by using optical sheets instead of epifluorescence exctitation.^10^ While both LCCT and HILO-fPT are capable of suspended phenotype cell 3D imaging, they require dielectrophoretic single-cell manipulation. Dielectrophoresis is an electrokinetic technique known to optimally operate at low conductivity buffers (∼10-100 mS m^−1^) to prevent Joule Heating,^11,12^ which can potentially damage the cell membrane, even at high field frequencies.^4^ Moreover, the serial approach to single-cell handling required in these micropatterned electrode-based techniques further complicates scalability.

At the heart of OPT is the ability to manipulate the 3D orientation of the imaged object relative to the optical axis of the system to generate projections. Common methods used to achieve controllable cell rotation and simultaneous OPT observation include microcapillary rotation^13^ and electric field induced cell electrorotation,^4,11^ which are effective but inherently low-throughput. Additionally, microcapillaries require cells to be embedded in cytotoxic, thixotropic index-matched gel, while in electrorotation, low conductivity buffers different from normal isotonic media are typically needed,^14^ thus limiting the biological assays that can be conducted during observation. In contrast, microfluidic cell rotation methods have also been explored, which provide less restrictive media conditions at higher throughput. For instance, flow-based approaches in microchannels have been used for quantitative phase imaging of cells based on hydrodynamic rotation,^15,16^ and acoustically-driven bubble arrays have demonstrated large-scale single-cell rotation compatible with tomographic reconstruction algorithms.^17,18^ Nonetheless, despite there being various microdevice-based cell rotation techniques (see ref. [19] for a recent comprehensive review), there remains the need for cell handling techniques that simultaneously provide: 1) parallel single-cell rotation suitable for 3D isotropic optical analysis of populations of suspended cells at the diffraction limit; 2) compatibility with fluorescence imaging modalities; and 3) an individualized microfluidic environment for live single-cells – particularly, unfixed cells susceptible to movement, such as immune cells.

Here, we present a passive microfluidic droplet trap array that leverages oil flow-induced interfacial shear to generate intra-droplet microvortices (Fig. 1a,b). These microvortices, in turn, mobilize encapsulated cells inside the water-in-oil droplets at an angular velocity determined by the exterior phase applied flow rate (Fig. 2). This strategy allows observation of cells encapsulated inside picoliter droplets of conventional, non-toxic isotonic buffer, and facilitates scalable, arrayed OPT acquisition by the simultaneous spinning of cells (see Fig. 1c and Movie S1). Our Arrayed-Droplet OPT (ADOPT) imaging modality enabled the isotropic 3D imaging of live myeloid and lymphoid cells in suspension and at the optical diffraction limit (x,y,z resolutions ∼216 nm with λ=461 nm, NA=1.3). This allowed morphometric analysis of K562 cells, as well as naïve and activated healthy-donor derived T cells, which are prone to motion in their suspended state. Furthermore, ADOPT demonstrated intracellular and fluorescence surface imaging of cells using a simple, camera-based epifluorescence microscopy set-up. Owing to this camera-based, single-cell continuous signal recording strategy, we were able to perform pixel-to-pixel image restoration through temporal filtering of photon influx events damaged by noise. This approach significantly improved noise characteristics of the standard CMOS detector here used, especially at low SNR ratios. Moreover, ADOPT of naïve and activated primary T cells facilitated nuclear content morphometry, which identified six distinct nuclear shape categories across immunological activation states over the course of seven days. Collectively, our findings present a novel droplet microfluidics framework that enables high-resolution, 3D isotropic analysis of non-adherent cells, opening avenues for population-based investigations into their dynamic intracellular and topographic features.

**Fig. 1.**
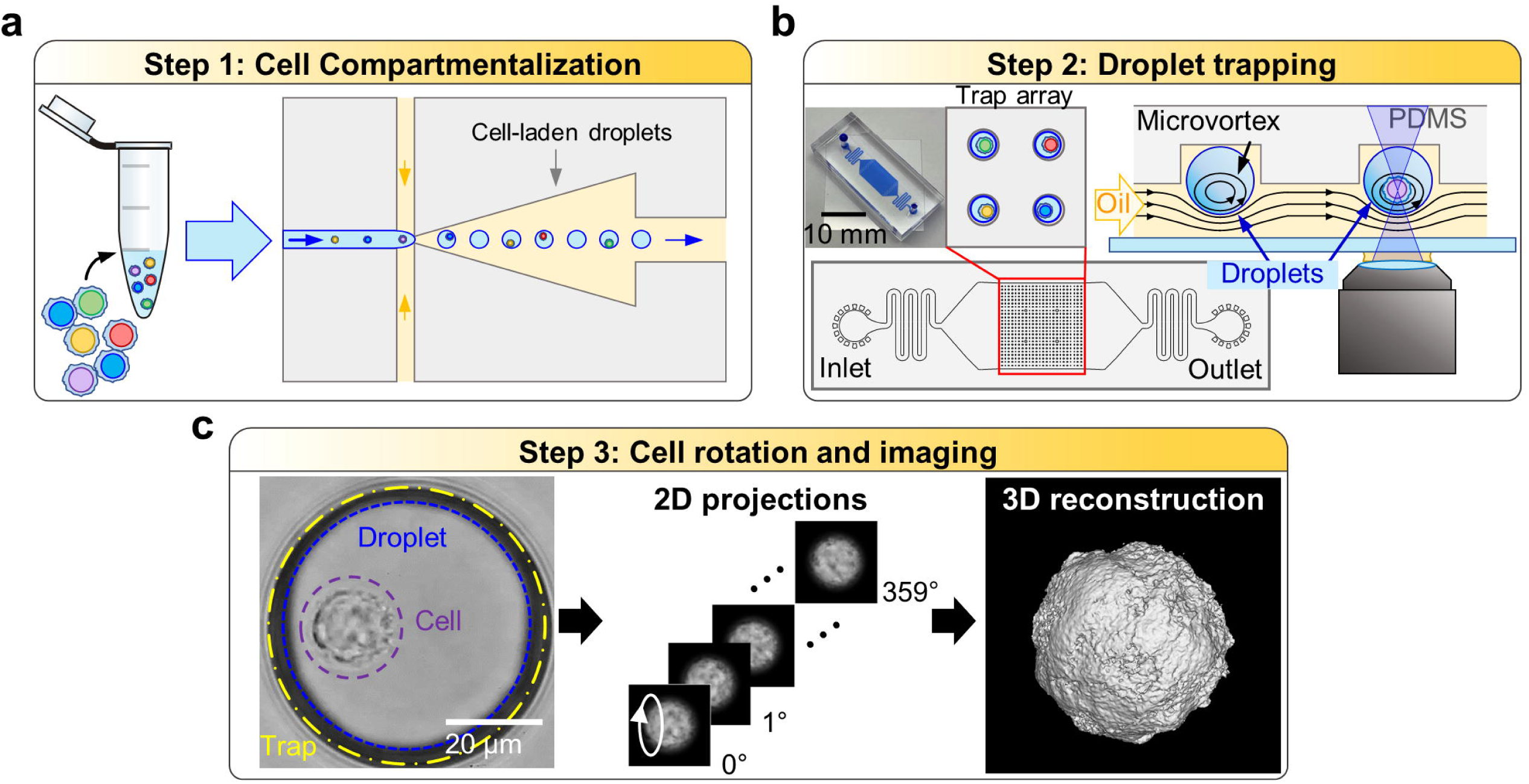
Workflow for 3D isotropic imaging of live suspension cells via ADOPT. a) A structurally heterogeneous population of cells is first compartmentalized into water-in-oil droplets. b) Cell-laden droplets are then loaded into a PDMS microfluidic trap array (24×24). Each trap consists of an inverted, circular microwell that can trap exactly one cell-laden droplet. Continuous oil perfusion transfers shear-stress to the inside of droplets, generating recirculation microvortices that drive cell self-rotation. c) In the last step, time-lapse frame acquisition of the rotating cells allows their full 360° observation, which can be used to approximate their 3D structure through PSF-informed OPT reconstruction algorithms.

**Fig. 2.**
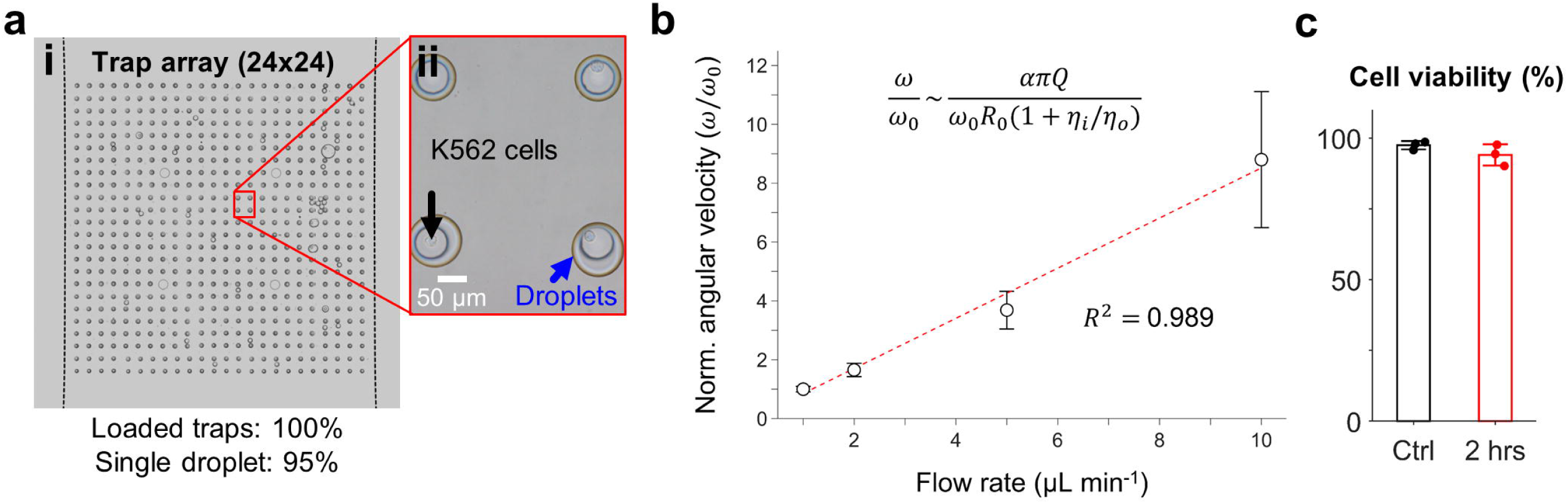
Flow rate induced cell rotation in microfluidic droplets. a-i) Microfluidic trap array image obtained via image stitching, showcasing 100% loaded traps with 95% single droplet occupancy. a-ii) Blow-up of droplet traps with encapsulated K562 cells. b) Normalized angular velocity of encapsulated cells as a function of external oil flow rate. c) K562 cell viability inside droplets when subjected to 2 hrs of continuous oil flow rate (1 µL min^-1^).

## Materials and Methods

### Microfluidic chip fabrication

Arrays of droplet traps, made from polydimethylsiloxane (PDMS), were manufactured through a two-layer standard soft-lithography process. For the photolithography steps, a first 40 µm thick layer of SU-8 3025 (Kayaku, MA, USA) was spin coated on a clean Si wafer (100) at 500 rpm for 10 s at an acceleration of 100 rpm s^-1^, followed by 2000 rpm for 30 s at an acceleration of 300 rpm s^-1^. This was followed by soft baking at 95°C for 15 min, UV exposure (365 nm) at a dose of 150 mJ cm^-2^, and a two-step post exposure bake (PEB) of 1 min at 65°C and 3 min at 95°C. A second 30 µm thick layer was thereafter coated at 2500 rpm following the aforementioned acceleration procedure. Soft-bake was then carried out at 95°C for 10 min, a second UV exposure at ∼125 mJ cm^-2^, and the previously mentioned PEB steps. The SU-8 material was developed for 7 min, with the last minute aided by sonication to completely etch the inverted microwell circular structures (traps).

After photolithography, Sylgard 184 monomers and silicone elastomer curing agent (Dow Corning, USA) were mixed at a 10:1 ratio and cast into the SU-8 microfluidic master structures. The liquid mixture was degassed for 1-2 hours in a desiccator and then allowed to partially cure overnight at room temperature. Subsequently, the mixture was heated to 65°C for 24-48 hours in a convection oven. The solidified PDMS material was removed from the molds, hole-punched, and any debris was cleared using scotch tape. Finally, the PDMS device was plasma treated (Harrick Plasma, USA) for 1 min and bonded to clean, thin glass slides (170 µm thick).

### Droplet encapsulation of cells

A previously described microfluidic device^20^ with 40 µm in height was employed for the generation of droplets containing suspension cells. Specifically, the continuous phase consisted of 2% (w/v) 008- FluoroSurfactant in HFE 7500 oil (Ran Biotechnologies, USA), while the dispersed phase comprised 1x PBS, 16% Optiprep (STEMCELL Technologies, Canada) along with the suspended cells. Cells were washed and prepared to a concentration of 10^7^ cells mL^-1^ in the dispersed phase. For encapsulated dead cell exclusion and viability analysis, propidium iodide solution (Biolegend, USA) was added to the dispersed phase at a concentration of 2 µL per 100 µL of cell solution (with 1×10^6^ cells mL^-1^), and incubated at 4°C for 15 min prior to imaging.

Picoliter-sized droplets were generated using two Flow EZ pressure control modules (LU-FEZ-7000, Fluigent, USA). First, microdevices were primed with the pressure pump delivering the continuous phase at a pressure of 115 mbar. Once devices were primed, the second module was set to 120-140 mbar, which resulted in squeezed cell-laden droplets with a diameter of 65-71 µm, as seen from 2D images from a conventional widefield inverted microscope at 10x magnification (Eclipse TE2000-S, Nikon, USA). Specifically, smaller droplets (65 µm diameter from circular view) were optimal for imaging naïve T cells (5-7 µm), whereas bigger droplets (71 µm) performed best for activated T cells (6-12 µm) and K562 cells (10-20 µm). Lastly, droplets were collected for 30 min and subsequently stored in 1.5 mL Eppendorf tubes (Eppendorf, Germany).

### Droplet entrapment using inverted microwell array

Prior to the droplet loading experiment, devices were vacuumed for 5 min to prevent the formation of any bubbles during sample input, and thereafter primed with continuous phase oil. After droplet generation and trap device priming, 10 µL of cell-laden droplets were pipetted into the inlet reservoir of the microfluidic trap array device. A pipette tip (200 µL) inserted into the inlet of the device acted as a reservoir for droplets. Continuous phase oil was then perfused into the chip using a syringe pump in withdrawal mode at 4 µL min^-1^ (Pump 11 Elite, Harvard Apparatus, USA), which eventually resulted in droplets loading into the array. Once all traps were occupied by droplets, the flow rate was set to 1-2 µL min^-1^ to perform cell rotation for 3D live cell imaging. Cell rotation rates were also characterized using a custom-made routine (MATLAB, USA) as a function of the applied continuous phase flow rate (1-10 µL min^-1^).

### Arrayed-droplet optical projection tomography (ADOPT)

After droplet loading into microfluidic traps, an oil flow rate of 1-2 µLmin^-1^ was used to produce rotation rates of 4.2-8.4 rpm. This allowed acquisition of N=800-1200 slices per cell at a 6.67 ms exposure (150 fps) which were obtained using an Olympus UPlanFL N 100x oil immersion objective (NA=1.3) mounted on an Olympus IX51 epifluorescence microscope (Olympus, USA). Our OPT routine is based on a previously reported direct inversion algorithm which helped reduce noise from out-of-focus light.^21^ Furthermore, monochrome 204×204 px 12-bit frames were collected using a CS505MU Kiralux 5.0 MP monochrome CMOS camera (Thorlabs, USA) with a 3×3 binning scheme to increase SNR. These settings resulted in an xy image resolution of 79 nm px^-1^.

Prior to building the 3D volumetric reconstructions from CMOS footage, pixel-to-pixel image restoration was carried out by performing signal filtering. Briefly, for each set of time-lapse frames containing a full rotation of a cell, the time-dependent signal of each pixel was filtered using a third-order, zero-phase Butterworth digital filter with a defined cut-off frequency, *f*_*c*_ (1 – 50 Hz). After performing SNR optimization (Fig. 4f), *f*_*c*_ = 5 Hz was chosen to filter all collected time-lapse frames in this study, unless otherwise specified.

To visualize cell surface topography, K562 cells and T cells were stained with anti-CD45 AlexaFluor 488 (HI30, Invitrogen, US). Intracellular imaging of cell nuclei was performed using NucBlue™ (Hoescht 33342 dye, Invitrogen, US) by following the manufacturer’s instructions. For T cell activation experiments, the nuclear content morphometry of 100 cells was assessed on days 0, 3 and 7, and classified according to its corresponding morphology (spherical, bean-shaped, ellipsoid, multi-lobed, dispersed, or amorphous).

### Cell culture

K562 cells obtained from the American Type Culture Collection (ATCC, USA) were cultured in IMDM medium (Gibco, USA) supplemented with 10% heat-inactivated fetal bovine serum (HI-FBS; Gibco, USA) and 100 U mL-1 penicillin–streptomycin (Gibco, USA). Cells were consistently maintained in a humidified incubator at 37°C with 5% CO_2_ and passaged every 2-3 days using standard procedures. Cell encapsulation in droplets and trapping experiments were conducted on the same day that cells were subcultured.

### Primary T cell isolation, activation, and expansion

Healthy donor blood samples were procured from the Institute for Clinical and Translational Science (ICTS) at the University of California, Irvine. Within less than 3 hours from blood collection, primary human T cells were obtained via standard immunomagnetic bead-based negative selection. Direct Human T cells isolation kits (EasySep™, STEMCELL Technologies, USA) were used according to the manufacturer’s instructions.

Primary human T cells underwent activation and expansion using standard bead-based protocols. Initially, freshly isolated T cells were seeded at a concentration of 0.2-0.5×10^6^ cells mL^-1^ in 24-well plates with 1 mL of RPMI 1640 medium, ATCC modification (Gibco, USA). The medium was supplemented with 10% HI-FBS (Gibco, USA), 2 mM L-glutamine, 1 mM sodium pyruvate, 0.1 mM non-essential amino acids, 10 mM HEPES, and 100 U mL^-1^ of penicillin-streptomycin (Gibco, USA). To activate and expand T cells, media-washed human activator CD3/CD28 Dynabeads (Gibco, USA) were added at a 1:1 cell-to-bead ratio, and the media further supplemented with 30 U mL^-1^ of human recombinant IL-2 protein (STEMCELL Technologies, USA). T cells were incubated in a humidified incubator with 5% CO_2_ at 37°C and were split to 0.5×10^6^ cells mL^-1^ when their concentration surpassed 2.5×10^6^ cells per well or when medium became depleted (turning yellow in color). Expansion fold numbers were recorded on days 0, 3 and 7.

### T cell immunophenotyping

Human T cells were concurrently stained with anti-CD4 Pacific Blue (OKT-4, Biolegend, USA), anti-CD8a APC (RPA-T8, Invitrogen, USA), and either anti-CD25 FITC (BC-69, Biolegend, USA) or anti-PD-1 FITC (MIH4, Invitrogen, USA) to determine their levels of activation during expansion experiments (days 0-7). Standard washing and incubation procedures for flow cytometry were followed from Biolegend. Dye 7-AAD (Biolegend, USA) was used for dead cell exclusion from the analysis. A NovoCyte 3000 flow cytometer (ACEA Biosciences, CA, USA) was used to analyze the cell suspension samples. Single-color stained cells were used to determine compensation levels, whereas positive and negative control samples were used to determine gating levels for quantification.

### Unfixed T cell drift tracking

The movement of T cells inside 24-well plates without any treatment or fixing was tracked to quantify drift in their suspended phenotype. Conventional brightfield images were collected in widefield mode using an Olympus IX51 microscope equipped with a 10x air objective. Videos were collected at 50 fps, with 1224×1080 resolution at 2×2 binning using a CS505MU Kiralux 5.0 MP monochrome CMOS camera (Thorlabs, USA). A multi-object tracking algorithm built in Matlab was used to create a population metric of the time-lapse drift of cells (see Fig. 6e).^22^ The total cell displacement, 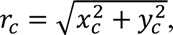 where (*x_c_*, *y_c_*) was the position of cells in each individual frame, was calculated for a total of *N* = 94 cells.

### Immunofluorescent confocal imaging

Confocal imaging of fixed and unfixed human T cells on days 0, 3 and 7 was performed using an Olympus Fluoview FV3000RS CLSM. Briefly, cells were simultaneously stained with Hoescht 33342 dye and anti-CD45-AlexaFluor 488 to visualize surface and intracellular features, and excited using a one-way sequential line scan at 350 nm and 488 nm with a pinhole size of 167 µm and a scan speed of 2 µs px^-1^. In addition, non-activated T cells were imaged for control purposes. Cells were observed using an Olympus UPLSAPO 40x silicone oil immersion objective (NA=1.25). The obtained pixel resolutions were 78 nm px^-1^ in the xy direction, and 200 nm px^-1^ in the z direction. A z-step size of 200 nm was used to generate the 3D reconstructions of live fixed and unfixed T cells. With the described settings, calculated optical lateral resolutions were 201 nm, and 1.003 µm for the axial direction.

## Results

### Microvortex-driven cell rotation using droplet trap arrays

The ADOPT framework consists of three main steps: 1) compartmentalization of cells into droplets (Fig. 1a); 2) arrayed-droplet trapping via passive hydrodynamics and intra-droplet microvortex generation (Fig. 1b); and 3) large-scale single-cell 360° visualization for OPT 3D reconstruction (Fig. 1c). Cells compartmentalized with a previously reported device were prepared for loading into the droplet trap arrays.^20^ We found that as little as 80-100 µL of cell suspension was sufficient to prepare cell-laden droplets. Similarly, in the second step, we determined that 10 µL of water-in-oil, cell-laden droplets were sufficient to operate the microfluidic trap array and generate microvortex-driven 3D reconstructions.

Our microfluidic droplet trap device features a two-layer flow-through PDMS channel, with the first layer measuring 40 µm in height and the second layer, situated on top, measuring 30 µm in height (see device side-view in Fig. 1b). The top layer incorporates a 24×24 inverted circular microwell array which was designed to hold exactly one droplet per trap. The water-in-oil, cell-laden droplets can be mobilized into the passive hydrodynamic traps by perfusing the continuous phase oil at a constant flow rate. We selected droplets to have 65-71 µm in diameter (as seen from a top view), which after being loaded into the device were slightly squeezed into a pancake shape in the depth direction perpendicular to the flow. Thus, as droplets approached a circular microwell, they were captured by a combination of buoyancy (HFE75000 oil density of 1614 kg m^-3^) and surface energy minimization.^23^ As seen in Fig. 2a-i, the microfluidic trap array can be loaded with 100% droplet occupancy efficiency at 95% single droplet trapping rate.

To characterize the single-cell rotation dynamics, droplets were loaded with K562 suspension cells (Fig. 2a-ii). The 12-bit video output of a CMOS camera in digital numbers (DN) was monitored over the region of interest (ROI) of a single-cell per frame and was used to obtain the frequency at which cells rotated via Fourier analysis. We determined that external oil flow rate can be used to increase the velocity magnitude of microvortices inside droplets, which in turn modulates the aforementioned rotation frequency (or equivalently, cell angular velocity). As seen in Movies S1 and S2, applying a 2 and 10 µL min^-1^ oil flow rate resulted in significant angular velocity increase. Furthermore, Fig. 2b depicts how a ten-fold increase in flow rate (1 to 10 µL min^-1^) results in a roughly ∼9-fold angular velocity increment. To explain the approximately linear relationship in Fig. 2b (*R*^2^=0.989), we derived the recirculation timescale (τ) of an infinitesimally small particle following the closed-loop vortex streamlines of a Hadamard-Rybczynski velocity field and applied it to estimate the angular velocity timescale. We made this approximation based on the observation that recirculation patterns within stationary droplets exposed to external uniform flows generate similar fields. This assertion, however, does not account for the complex effects of trap geometry on fluid flow. Following this argument, we found τ is given by τ∼2*R*_0_/*U*_0_(1+*η_i_ /η_o_*), where *R*_0_ is the droplet radius, *U*_0_ the bulk flow velocity and *η_i_ /η_o_* the ratio of inner and outer phase viscosities. Assuming the cell rotation period is *T*∼*τ*, and *U*_0_ is proportional to the applied flow rate (*U*_0_*∼αQ*), ω becomes:

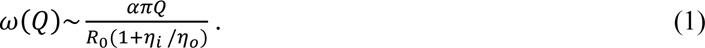

We further tested the effects of droplet-induced cell rotation on cell viability. It was determined that 2 hrs of continuous rotation driven at 1 µL min^-1^ resulted in a viability change from 97.5% to 94.0% (*p* = 0.1268, one-sided t-test).

### Imaging of cellular and subcellular 3D features in live non-adherent cells

ADOPT can be used to acquire immunofluorescent 3D volumetric data from single suspended cells with isotropic resolution. Specifically, we investigated its performance to quantify the presence of CD45 markers on the surface of K562 cells. Figure 3a portrays the maximum intensity projection (MIP) of the fluorescent signal, while Fig. 3b takes an isosurface rendering of that 3D reconstruction using an isovalue of 0.4 with respect to the normalized fluorescence intensity in arbitrary units. These reconstructions evince the isotropic resolution available to OPT imaging methods. Similarly, the topography of smaller cells, such as primary T cells (activated and non-activated) can be imaged with isotropic resolution at the diffraction limit, as estimated by the Rayleight criterion (0.61 λ/NA = 244 nm for this fluorophore). For example, a rotation slice from a CD25+ T cell labelled with CD45 surface markers is shown in Fig. 3c, along with its corresponding isotropic surface topographic reconstruction in Fig. 3d.

**Fig. 3.**
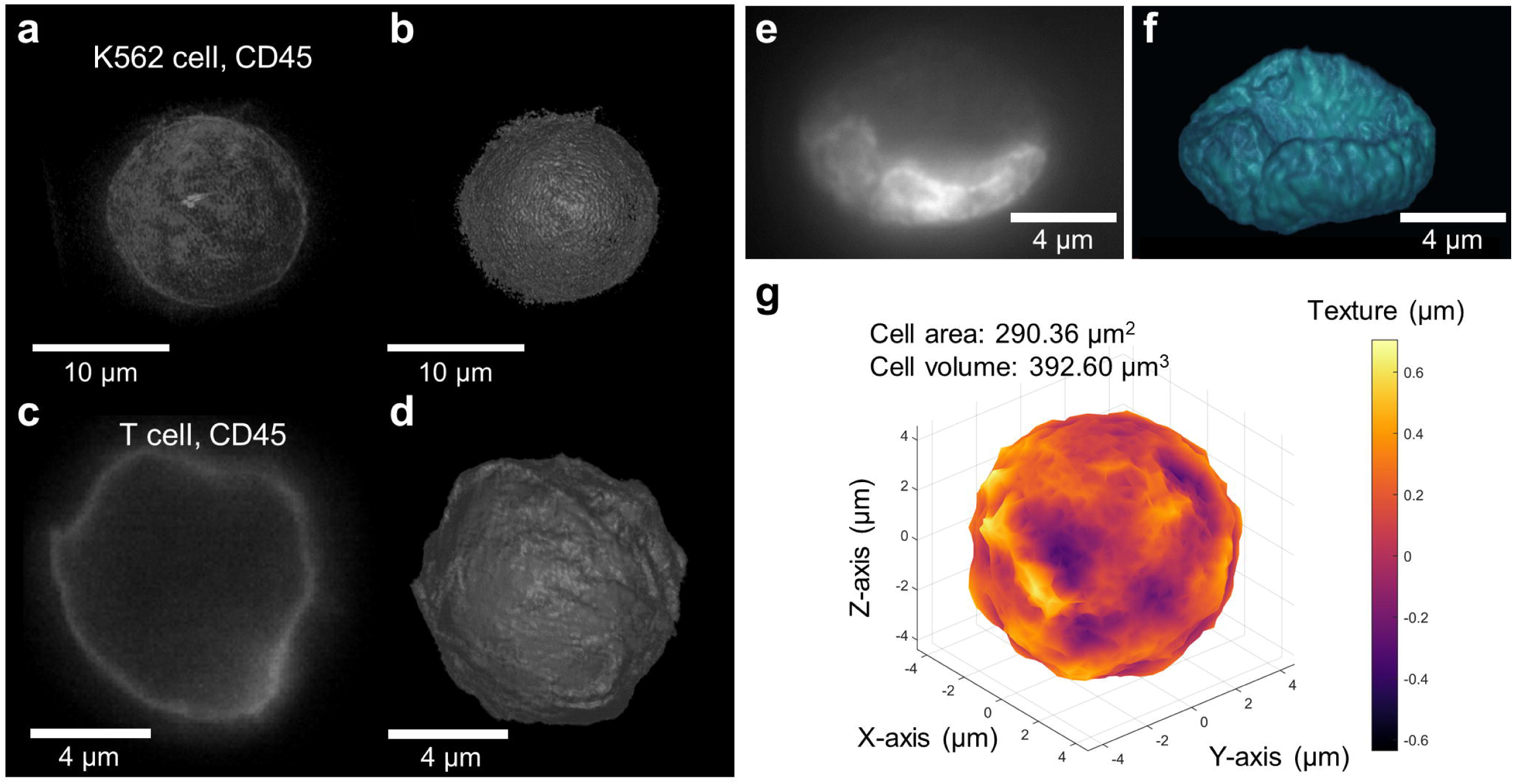
Three-dimensional live cell topographical and intracellular imaging with isotropic resolution via ADOPT. a,b) Suspension cells (K562 cell line) stained with anti-CD45 - AlexaFluor 488, visualized via its MIP (a), and isosurface volumetric rendering (b). c,d) Activated T cell (CD25+), marked with anti-CD45-AlexaFluor 488 for visualization purposes. c) A 2D slice taken while acquiring the required 360° for 3D reconstruction via OPT. An isosurface of the CD45 fluorescence level is shown in (d). e) 2D image of the stained nucleus (Hoescht dye 33342) of a K562 cell. f) 3D reconstruction of the nuclear content distribution of the nucleus in (e). g) 3D isosurface of a K562 cell, detailing quantitative data of its texture with a heatmap, as well as its volume and surface area.

We further demonstrated that ADOPT can readily reconstruct intracellular structures like the DNA content of the nucleus. Diffraction limited images of the nuclear content in live K562 cells, made visible through Hoescht 33342 staining, are displayed in Fig. 3e. Collection of N=800 slices at varying angles (0.45°/slice, 150 fps, raw footage in Movie S3) were used to generate the 3D volumetric rendering in Fig. 3f (Movie S4), which features an estimated isotropic resolution of 216 nm.

For all 3D surface models shown, morphometry can be used to retrieve single-cell topographic, quantifiable data. The surface model in Fig. 3g presents the texture map of a K562 cell surface, where the color of each surface patch represents how far it is from the 3D model average radius. This measurement is therefore closely related to the cell surface roughness and can potentially be used to compare single-cell metrics in a population manner for textural variations. Other single-cell morphometry parameters that can be calculated from our workflow include cell area, volume or other morphological descriptor that offers enough discriminatory power, such as sphericity.^24^

To improve the quality of reconstructions, we illustrated in Fig. 4 how the SNR of the collected time-lapse single-cell rotation frames can be greatly enhanced by using a pixel-to-pixel image restoration strategy. Considering that each set of frames containing the rotation of a cell is continually captured with a camera-based system (CMOS), the time-dependent photon influx information of each pixel can be restored by eliminating high-frequency components in the readout. This filtering process can be seen in Fig. 4a-d, where the raw signal from a CMOS camera (Fig. 4a) is restored by applying a third-order, zero-phase low-pass Butterworth digital filter (Fig. 4b). In these figures, the nuclear content of a rotating K562 cell was visualized via live-cell staining. In Fig. 4e, the application of the described filter is demonstrated, showcasing the denoising of time-lapse information from a single pixel in Fig. 4c, collected over the full cell rotation period (*t* = *T*). Furthermore, blow-up colormap plots in Fig. 4b,d demonstrate the improved image quality achieved via filtering, which enhanced the visualization of the coarser chromatin boundaries present in the nucleus. This is further evinced by line-profiles of the restored micrographs, which allowed estimation of the distance between these chromatin boundaries (∼0.5 µm).

**Fig. 4.**
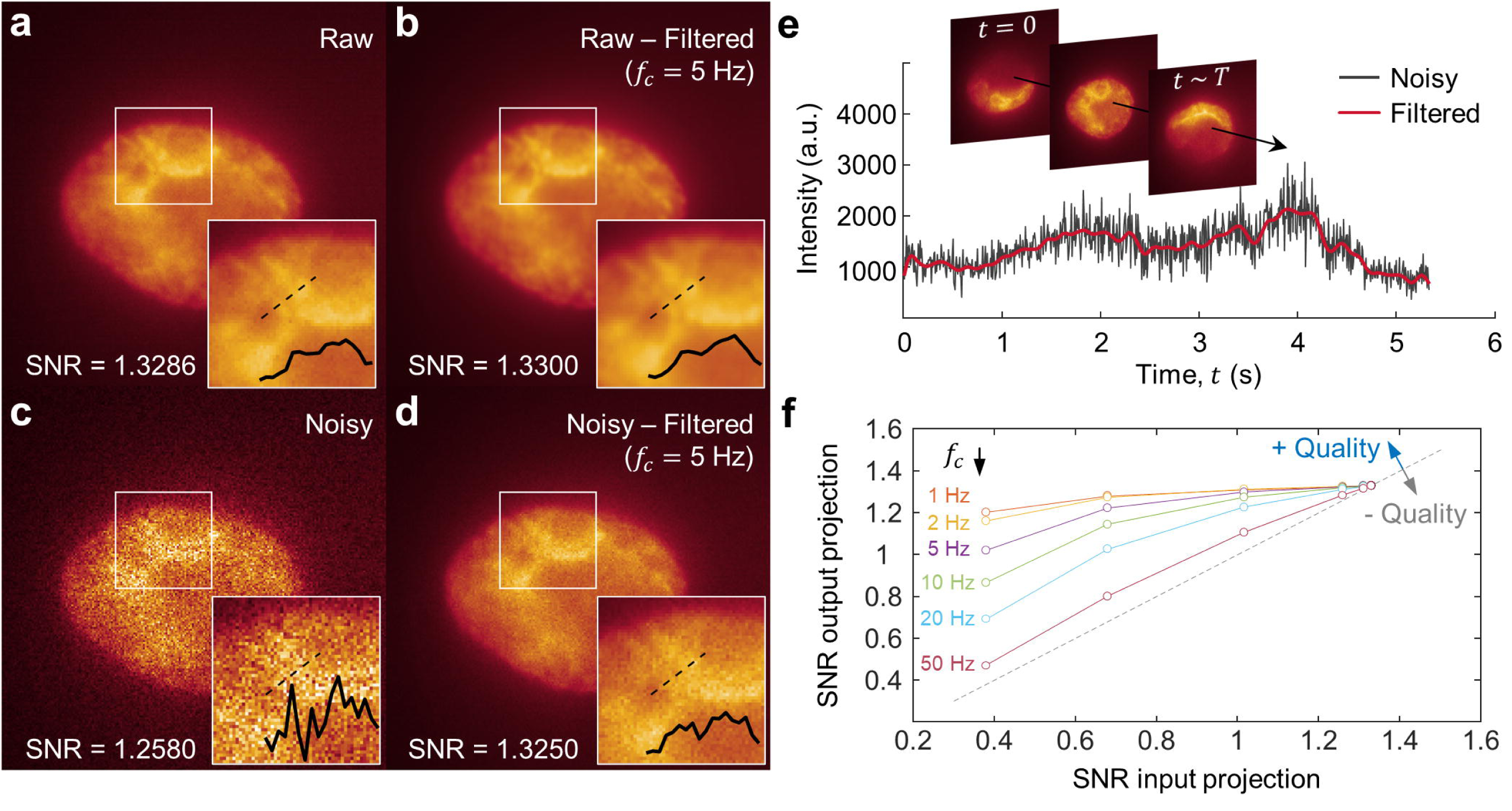
Pixel-to-pixel denoising of 2D projections obtained via single-cell rotation. a) 100X raw image (i.e. projection) of a K562 cell nucleus stained with Hoescht 33342 dye. Insets show a blow-up of the boundary between two chromatin-rich structures and their corresponding line-profiles. b) Denoised image from a), obtained by filtering the time-lapse signal of each pixel with a Butterworth filter (cut-off frequency, *f*_*c*_ = 5 Hz). c) Image in a) with added Gaussian noise (SNR = 1.2580). d) Denoised image from c), following same method as in b). e) Time-dependent signal coming from a single pixel (time evolution schematically emphasized by arrow). The plot demonstrates the use of a Butterworth filter to eliminate high-frequency components (low-pass filter). f) Image quality improvement (measured as output SNR) as a function of the input noise level (SNR) and cut-off frequency *f*_*c*_. Dashed line represents no net improvement (one-to-one SNR of input and output).

Figure 4c,d further provides evidence that highly distorted images (SNR = 1.2580) can be restored to SNR levels that allow the distinction coarser chromatin features. We validated the performance of our filtering algorithm by providing images with known SNR levels and found that signal improvement was largely dependent on the starting SNR values, as well as the low-pass filter cut-off frequency (*f*_*c*_). By studying the effect of varying *f*_*c*_ (Fig. 4f), we identified 5 Hz as providing a balance between enhanced SNR rates, while preserving high frequency components in the image coming from the actual, time-dependent light readout.

### ADOPT performance vs. optical sectioning techniques

#### Optical sectioning SNR

We compared the SNR characteristics of 3D images collected via CLSM and ADOPT. While CLSM remains a major workhorse in biomedical optical microscopy for its out-of-focus light rejection capabilities (e.g. Fig. 5a), it is susceptible to high depth-dependent degradation.^5^ We observed this artifact was present in z-stacked images of substrate-bound T cells labelled for surface and intracellular markers, as seen in Fig. 5b-d. In contrast, T cells imaged with the ADOPT strategy were not visibly damaged by sectioning artifacts, as seen in Fig. 5e,f. To quantify the degree to which the optical signal was degraded per collected slice, we computed its corresponding SNR, given by the ratio of signal average intensity (*μ*) and standard deviation (*σ*) of each slice. After normalizing for optical intensity, we observed ∼3.2-fold decrease in SNR in confocal z-stacks when comparing the initial and final z-stack levels to mid-range slices (Fig. 5g). On the other hand, the cell-rotation slicing produces marginal changes (∼1.2-fold) in SNR. Lastly, we note that the volumetric reconstructions collected by optical sectioning had visible lower axial resolution owing to the PSF anisotropy. This observation was consistent with the calculated lateral and axial optical resolutions for the CLSM set up, which were 201 nm, and 1.003 µm, respectively.

**Fig. 5.**
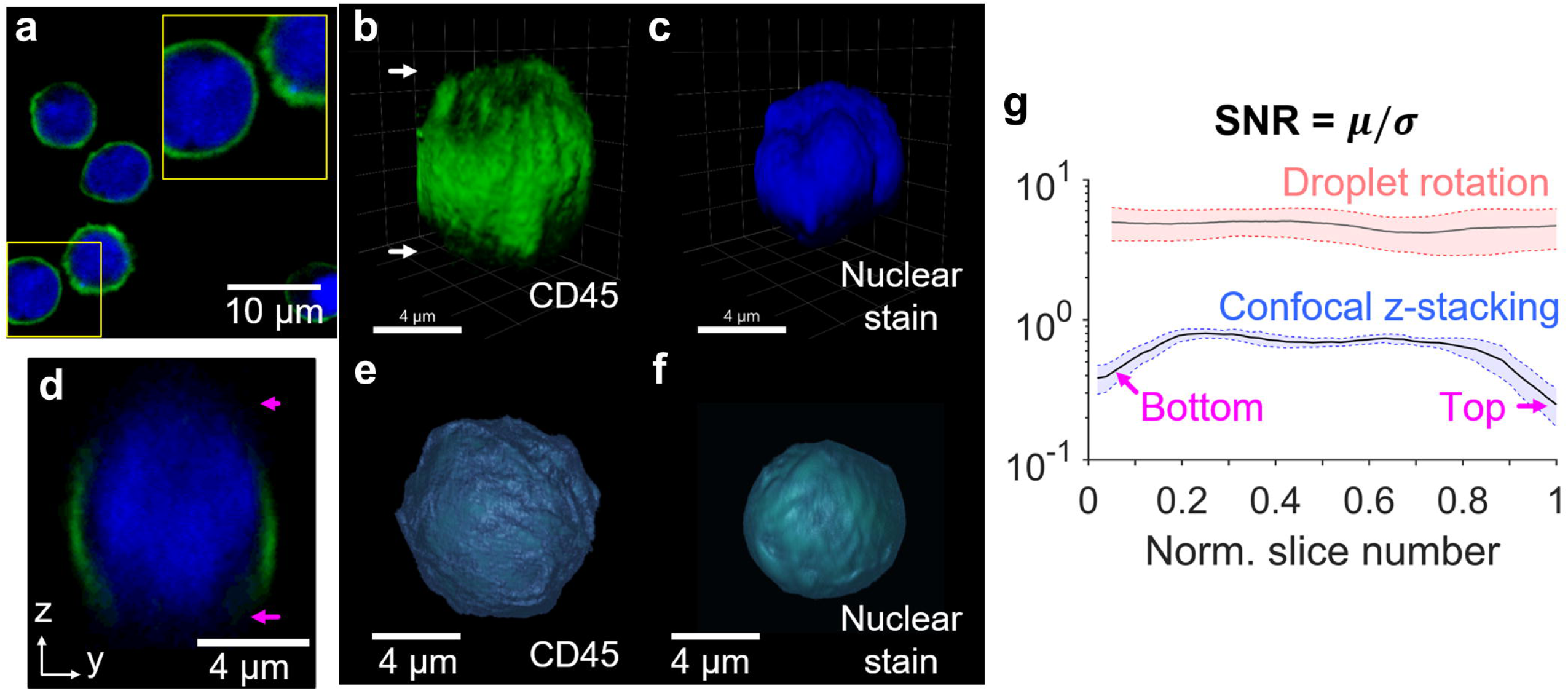
a) Confocal optical section of naïve T cells bound to the substrate, with their surface marked with anti-CD45-AlexaFluor 488 and their nucleus with Hoescht 33342 dye. Blow-up shows the distribution of nucleus with marked regions resembling lobes. b,c) Z-stacking images of the surface (b) and nucleus (c) of T cells, showing the top and bottom artifacts in the reconstruction, as well as the reduced axial resolution. d) An xy-plane cut further evinces the top and bottom optical sectioning artifacts. e,f) ADOPT reconstructions of the surface and nucleus of an activated T cell. g) Comparison of the SNR characteristics of ADOPT vs. CLSM as a function of the slice number (normalized by N = 61 slices for confocal and N = 500 slices for the droplet rotation case). AlexaFluor 488 fluorophore was analyzed for (g) by triplicate in each case, and the frame grayscale values were normalized by the peak intensities in each case. Boundaries represent the average SNR values ± the standard error of the mean (s.e.m.).

#### Drift artifacts of suspension cells

Observing non-fixed immune suspension cells by z-stacking methods presents a challenge, considering the temporal resolution in techniques that perform pixel-wise scanning of biological samples. The presently tested CLSM equipment operated with pixel scanning dwell times of ∼2 µs px^-1^, which even when evaluating low resolution ROIs (256×256 to 512×512 for single-cell observation), brought the slice acquisition time close to the minutes scale for N=100 stacks (∼20 µm in z-direction). The effects of these prolonged capture times can be seen in the 3D live cell images and slice visualizations of Fig. 6a-d, where non-fixed T cells showed signs of both rotational and translational drift artifacts. Moreover, by performing multi-cell centroid tracking in time-lapse videos, we quantified the average drift of N=94 unfixed cells and plotted the results in Fig. 6e. Interestingly, the observed tendency was that T cells drift monotonically overtime, reaching close to 0.8±0.4 µm displacements in ∼3.8 seconds. Conversely, cells that spin about their rotation axis driven by microvortices experience a deterministic, value-bounded average displacement (0.3±0.1 µm) that is periodic in nature.

**Fig. 6.**
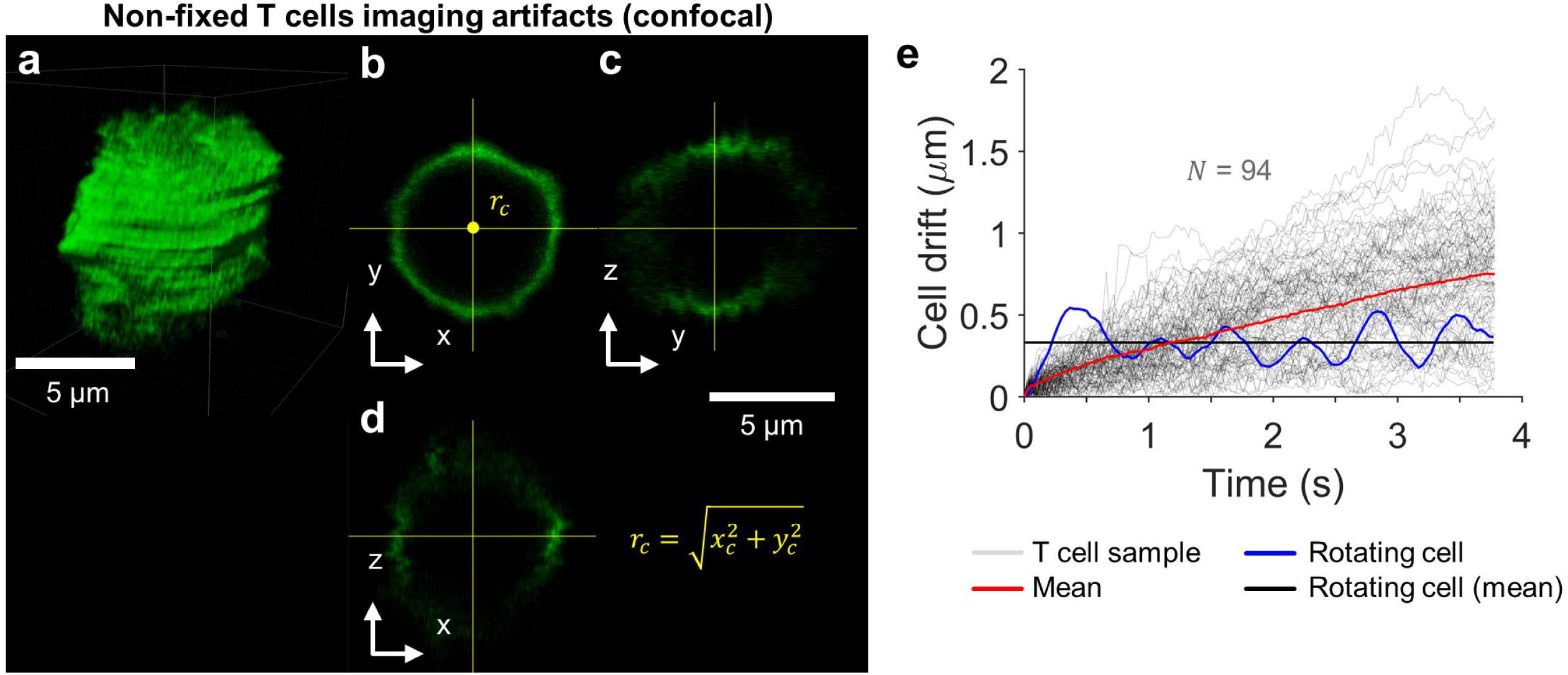
a) Translation and rotation motion artifacts of unfixed T cells, as imaged by a CLSM system (fluorophore: anti-CD45-AlexaFluor 488). b) A single xy-plane slice of the T cell in (a), showing its central position for tracking. c,d) Different plane cuts illustrating motion artifacts. (e) Cell drift as a function of time, traced for N = 94 T cells. The calculated drift (in the xy-plane) of the centroid of a rotating cell inside a droplet. Average values are highlighted to evince motion trends.

### Primary T cell activation phenotype and nuclear morphometry

As previously shown, lymphocytes are particularly challenging to image without fixing protocols, and thus are a fitting candidate for single-cell, 3D live isotropic imaging with ADOPT. To get a general picture of the T cell activation phenotypic changes, T cells were isolated from healthy donor whole blood samples (N=3) using immunomagnetic negative selection. Cells were then imaged on days 0, 3 and 7 after their in-vitro dynabead-mediated activation. We obtained 2D brightfield (Fig. 7a) and 2D fluorescence optical sections (Fig. 7b) images of activated T cells via widefield and CLSM, respectively. In Fig. 7b, we noted that control T cells display nearly circular or slightly bean-shaped nuclear distribution, consistent with previous reports.^25^ T cells exhibited an elongated phenotype, which was particularly pronounced on day 3 following activation and stimulation with human recombinant IL-2. Morphological changes were further accompanied by an increased motility in T cells, as they surveyed the surface of well plates for contacting activator beads. Nuclear content in elongated T cells by day 3 was observed to decondense and acquire a multilobed or dispersed appearance, as seen in the CLSM fluorescence 2D sections of Fig. 7b.

**Fig. 7.**
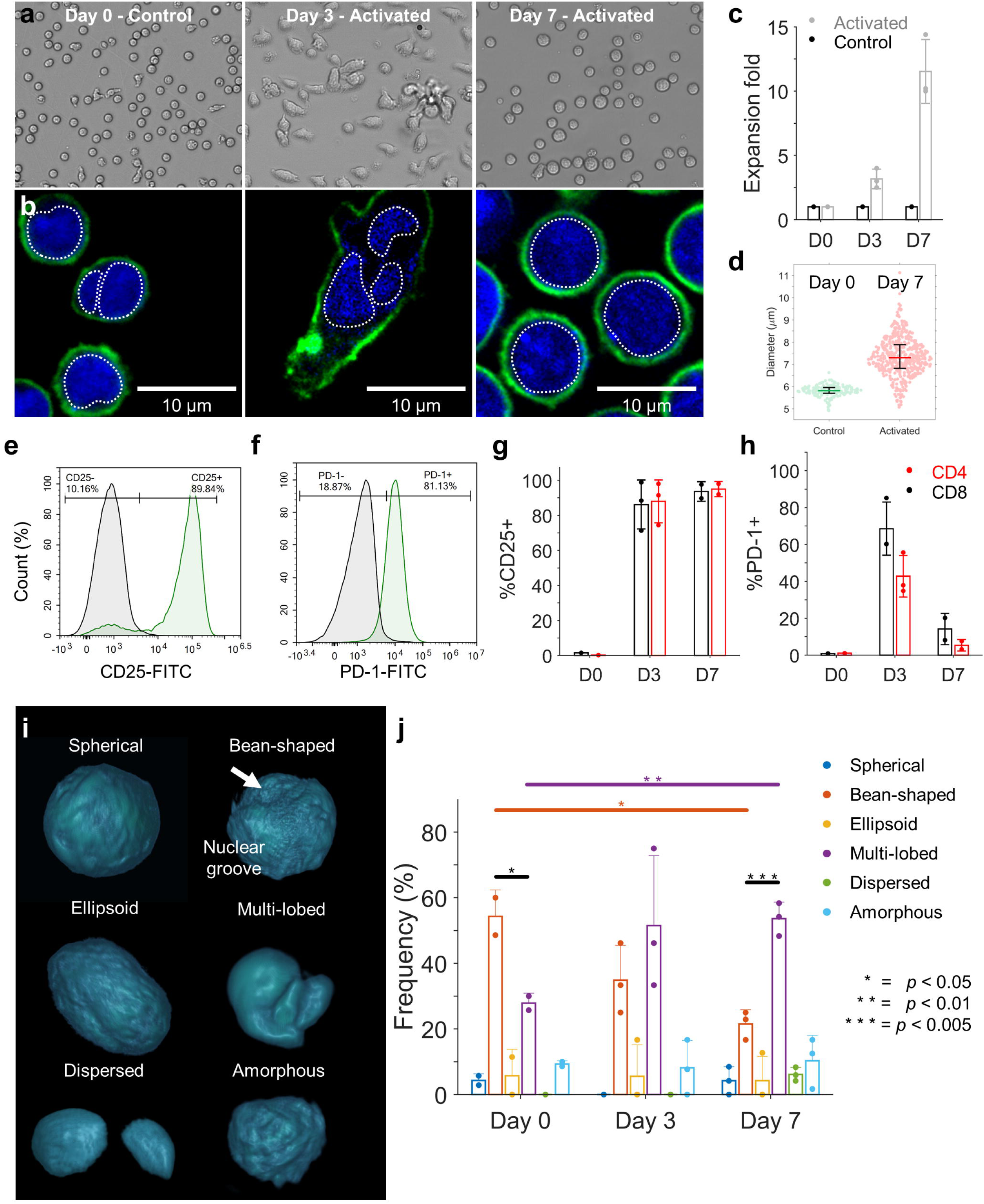
Primary T cell activation phenotype and nuclear morphometry. a) Brightfield images of T cells as a function of the number of days after Dynabead-mediated activation. b) Corresponding CLSM 2D optical sections, with cells stained for CD45 surface marker and nuclear content. c) Primary T cell expansion-fold as a function of elapsed days after activation, for N =3 healthy donors. d) Size distribution change from day 0 to 7. e,f) Flow cytometry analysis of the percent of CD25+ and PD-1+ pan T cells on day 3 post-activation, for donor 1. g,h) Percent of cells expressing CD25 and PD-1 markers for the N = 3 donors, plotted by days post-activation and classified by helper (CD4+) or cytotoxic (CD8+) phenotype. i) Activated T cell nuclear content morphologies identified via ADOPT 3D reconstructions. j) Distribution of T cell nuclear morphology phenotypes, shown as a function of days after activation (N = 3 donors, two-tailed t-test). Error bars throughout represent mean values ±standard deviation, except in d), where they represent quartile bounds.

Following human T cell activation (Fig. 7c), cells experienced size growth during expansion as quantified in Fig. 7d, from 5.8±0.3 µm to 7.3±0.9 µm, with a maximum size of nearly 11.1 µm by day 7. Other activation metrics were recorded, such as the percentage of T cells expressing CD25 and PD-1 markers (Fig. 7e-h), which were further classified by day and lymphocyte type (CD4+ or CD8+). The expanded T cells showed a sustained expression of CD25 markers for the duration of the experiment (7 days), as expected from bead-mediated activation protocols.^26^ Additionally, expression of PD-1 was highest by day 3 (∼65%) in cytotoxic CD8+ T cells. This result was consistent with the expected high activation levels in healthy cells, which at day 3 post-activation display peak values according to manufacturing standards.^27^

Next, we applied ADOPT to the study of nuclear morphological changes that primary human T cells underwent upon anti-CD3/CD28 immunomagnetic bead mediated activation. Owing to the 3D isotropic imaging capabilities of ADOPT, we were able to identify six distinct nuclear morphologies by visual inspection of the isosurface renderings. These consisted of spherical, bean-shaped, ellipsoid, multi-lobed, dispersed, and amorphous nuclear shapes (Fig. 7i). Furthermore, as seen in Fig.7j, analysis of N=100 activated T cells across seven days for N=3 donors illustrated an activation dependent distribution of these phenotypic nuclear morphologies. It was observed that most naïve T cells had a nearly circular bean-shape distribution, which after immunological activation markedly switched to a predominantly multi-lobed population. This was evinced by the significant change in the amounts of bean-shaped nuclei from day 0 to 7 (*p* < 0.05) and multi-lobed nuclei (*p* < 0.01) according to a two-sided t-test. Although other morphologies were qualitatively identified, their frequency was relatively low in comparison. Interestingly, while cells were analyzed in their suspended phenotype, the multi-lobed nature of the nuclear content identified in substrate-attached T cells Fig. 7b (day 3) was preserved. This suggests that the dispersed nucleus nature of motile, activated T cells might require additional time to return to its more condensed state.

## Discussion

Understanding T cell activation dynamics and their concomitant structural phenotypic changes is critical to the elucidating of their immune effector functions. Morphological changes, for instance, can include size growth and cytoskeletal rearrangement,^28^ which prepares cells for division and immunological synapse formation.^1^ Here, we have shown that T cell volumetric nuclear content undergoes significant reconfiguration in normal culturing conditions for immunological activation, and that these morphological changes persist after cells return to their suspended phenotype (Fig. 7i,j). We believe ADOPT is the first platform to demonstrate 3D immunofluorescent and isotropically-resolved reconstructions of fully suspended, live primary T cell surface and nuclear structures at the optical diffraction limit (Fig. 7i). These cells, being at the smaller size spectrum of leukocytes (5-7 µm), present a technical imaging challenge in their non-adherent state, drifting at rates capable of introducing significant artifacts, even on the timescale of seconds (0.2±0.1 µm s^-1^, Fig. 6). While attachment of T cells to coverslips using fibronectin produces excellent imaging results using high spatiotemporal resolution optical systems,^1^ it can also influence the microenvironment in which cells operate, as glycoproteins can affect cell movement and generate costimulatory signals that activates degranulation.^29,30^ These cues could therefore introduce extraneous integrin-mediated interactions that trigger cytoskeletal remodeling, cell motility or receptor signal modulation.^31^ In contrast, our device offers a scalable single-cell manipulation method without the need for surface treatments or cell fixing, making it, in principle, fully compatible with the specific liquid media required by the suspended (or trypsinized) cells. ADOPT therefore bridges a technological gap in live single-cell biomedical optics by leveraging the cell manipulation capabilities of intra-droplet microvortices, which were shown to produce periodic and stably bounded cell rotation (Fig. 6e).

Varied non-adherent cell nuclear structures were identified in this work (Fig. 3f and Fig. 7i), and the implications of such reconfigurations are potentially significant and warrant further investigation. Previous research has quantified different 3D nuclear shapes not immediately obvious in 2D cross-sections in normal, metastatic and fibrocystic cells.^24^ In that study, a phenotype deemed the “mushroom cap” nucleus shape was found as the predominantly morphology. This form closely resembles those observed in our experiments for K562 cells, which is a human immortalized myelogenous leukemia cell line (Fig. 3f). Furthermore, myeloid and cancerous cells tend to have bean-shaped, lobulated or segmented nuclei, configurations that have been posited to confer morphological flexibility.^25^ Indeed, the relationship between nuclear morphology and the coarser features of chromatin structural configuration has been associated with cancerous states in cells.^32^ Territories that chromosomes occupy are stratified, with transcriptionally silent domains being spatially located in the nuclear periphery, and content relocation varying occasionally with gene activation.^33^ Recognizing the crucial role of genome architecture in regulating transcription, DNA damage/repair, aging, and disease,^34^ ADOPT is poised to contribute to large-scale and multi-dimensional data-driven identification of morphometrically associated cell states and events.

Nuclear morphologies of T cells in different functional states have similarly been subjected to 3D volumetric analysis. While lymphocytes are generally recognized to have round, uniform and relatively rigid nuclear shapes that preserve genomic DNA integrity during circulation,^25^ events such as immunological activation can lead to actin-mediated elastic forces that deform their nucleus.^35^ T cells undergo constant shape changes in physiological conditions, resulting in actin-dependent nuclear elongation that correlates with gene expression, Erk and NF-κB signaling to the nucleus, and the expression of activation markers such as CD69.^36^ These activation-driven modifications can be understood in the context of immunological activation *in-vivo*. After activation, T cells exit lymph nodes, enter the bloodstream, and subsequently re-enter non-lymphoid organs, engaging in trans-endothelial migration to execute their effector functions.^37^ Trans-endothelial migration critically requires motor proteins, such as Myosin-IIA, to complete the movement of their rigid nucleus through endothelial junctions.^37^ Our nuclear distribution results in Fig. 7j would seem to support the hypothesis that a more dispersed nucleus content in day 3, with its concomitant CD25 and PD-1 expression increases, can prepare T cells for the functional challenge of undergoing heavy actin-driven phenotypic changes.

Other functional states in T cells associated with nuclear structure include the chromatin openness of specific genomic locations, such as the PD-1 genes. This chromatin openness has been shown to have predictive power for the effectiveness of anti-PD-1 immune checkpoint therapies in gastric cancer patients.^38^ Additionally, the study of other diseases, such as Nodular lymphocyte-predominant Hodgkin lymphoma (NLPHL) and T-cell/histiocyte-rich large B-cell lymphoma (THRLBCL), further emphasizes the importance of nuclear 3D analyses in activated T cells.^39^ Particularly, whereas 2D optical sections of T cell nuclei showed no significant enlargements, 3D volumetric measurements were necessary to show the significant differences between NLPHL and THRLBCL. We thereby anticipate that ADOPT quantification of multiparametric, immunofluorescence volumetric data of cells could not only potentially assess the functional state of T cells through nuclear measurements, but also help identify cell diseased states.

It should be mentioned that minute-frequency 3D imaging tomographic approaches, which quantify optical phase changes in single live cells,^18^ constitute important complementary technologies to ADOPT. While these methods possess non-isotropic resolutions (0.72 μm and 4.26 μm for the lateral and axial resolutions),^18^ they have shown scalability and label-free capacities, whereas ADOPT in its current rendition only demonstrates an epifluorescence widefield microscopy modality. In this sense, we believe the isotropic morphometry and specificity gained from chemical labelling are factors to be thoroughly considered in experimental design – especially in light of the single-cell analysis advantages that fluorescence activated sorting could bring when combined with droplet sorting preparative procedures.^40^ Other label-free optical methods have further demonstrated isotropically-resolved imaging based on tomographic molds for optical trapping,^41^ with resolution at the diffraction limit, but are incapable of suspended cell phenotypic imaging or multiplexed, chemically specific signal acquisition. Further limitations of ADOPT include its use of fluorescence OPT mode of acquisition, which is known to be prone to rotation artifacts far from the objects center of rotation.^42^ PSF aware reconstruction algorithms can be implemented to alleviate this issue, or a scanning focal plane scanning strategy,^21^ albeit at increased acquisition times. Despite these shortcomings, the simplicity of the wide-field and camera-based acquisition scheme utilized by ADOPT makes it an attractive 3D single-cell analysis tool. In its current form, the ADOPT components are not only cost-effective for most laboratory settings, but also compatible with pixel-to-pixel time-lapse denoising strategies (Fig. 4), which can be further enhanced by state-of-the-art, system-aware restoration techniques.^43^

ADOPT is further expected to be incorporated into the droplet microfluidic toolbox by providing single-cell 3D fluorescent marker spatial information that can be correlated with cell state and function. This approach can enable a wide range of applications relevant to T cell engineering, particularly in cancer immunotherapy research. Comprehensive immunophenotyping of T cell multifunctional responses and state progression during immunotherapy necessitates both the study of labelled surface antigens and secreted cytokines.^44,45^ Single-cell co-encapsulation in droplets with anti-cytokine detection probes has been shown to eliminate protein diffusive crosstalk typically seen in bulk assays^46,47^ – a strategy easily applicable to the ADOPT workflow, facilitating the correlation of 3D single-cell morphometric and secretomic information. In addition, mature technologies, such as single-cell droplet sorting create an opportunity for high-throughput, multiplexed analysis of specific cell subtypes, which could be readily integrated into the 3D morphological measurement workflow of our method. In essence, these advantages make ADOPT modular and customizable to upstream droplet preparative steps, thereby expanding the single-cell multidimensional analysis microfluidic toolbox for biomedical applications.

## Author contributions

B.C. B.: conceptualization, methodology, formal analysis, data curation, theoretical analysis, software, writing – original draft, review, editing and visualization. R.H.: methodology, formal analysis, data curation, writing – original draft, review, editing and visualization. X. L.: conceptualization, writing – review and editing. A. P. L.: conceptualization, methodology, resources, supervision, project administration, writing – review and editing.

## Conflicts of Interest

B.C. B., X. L., and A. P. L. are inventors of a US patent application related to this work.

## Acknowledgements

The authors would like to acknowledge support from the National Science Foundation and the industrial members of the Center for Advanced Design and Manufacturing of Integrated Microfluidics (NSF I-UCRC award number IIP-1841509).

