## Supplementary figures and images for "Three-dimensional Isotropic Imaging of Live Suspension Cells Enabled by Droplet Microvortices"

### Supplemental Movie S4

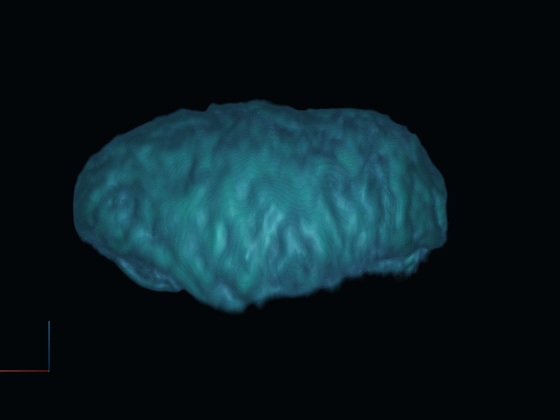
